# Evaluating performance of existing computational models in predicting CD8+ T cell pathogenic epitopes and cancer neoantigens

**DOI:** 10.1101/2020.12.25.424183

**Authors:** Paul R. Buckley, Chloe H. Lee, Ruichong Ma, Isaac Woodhouse, Jeongmin Woo, Vasily O Tsvetkov, Dmitrii S Shcherbinin, Agne Antanaviciute, Mikhail Shughay, Margarida Rei, Alison Simmons, Hashem Koohy

**Author notes:** These authors contributed equally (Paul R. Buckley, Chloe H. Lee).

## Abstract

T cell recognition of a cognate peptide-MHC complex (pMHC) presented on the surface of infected or malignant cells, is of utmost importance for mediating robust and long-term immune responses. Accurate predictions of cognate pMHC targets for T Cell Receptors (TCR) would greatly facilitate identification of vaccine targets for both pathogenic diseases as well as personalized cancer immunotherapies. Predicting immunogenic peptides therefore has been at the centre of intensive research for the past decades but has proven challenging. Although numerous models have been proposed, performance of these models has not been systematically evaluated and their success rate in predicting epitopes in the context of human pathology, has not been measured and compared. In this study, we evaluated the performance of several publicly available models, in identifying immunogenic CD8+ T cell targets in the context of pathogens and cancers. We found that for predicting immunogenic peptides from an emerging virus such as SARS-CoV-2, none of the models perform substantially better than random or offer considerable improvement beyond HLA ligand prediction. We also observed suboptimal performance for predicting cancer neoantigens. Through investigation of potential factors associated with ill performance of models, we highlight several data- and model-associated issues. In particular, we observed that cross-HLA variation in the distribution of immunogenic and non-immunogenic peptides in training data of the models seem to substantially confound the predictions. We additionally compared key parameters associated with immunogenicity between pathogenic peptides and cancer neoantigens and observed evidence for differences in the thresholds of binding affinity and stability, which suggested the need to modulate different features in identifying immunogenic pathogen vs. cancer peptides. Overall, we demonstrate that accurate and reliable prediction of immunogenic CD8+ T cell targets remains unsolved, thus we hope our work will guide users and model developers regarding potential pitfalls and unsettled questions in existing immunogenicity predictors.

## Introduction

The importance of being able to accurately predict CD8+ T cell targets i.e., immunogenic peptides, has never been clearer than during the ongoing COVID-19 pandemic era. It is also central in devising personalised vaccines for various cancers.

An efficient antigen-specific CD8+ T cell response to exogenous pathogens or endogenous threats relies on tightly regulated processing and presentation of antigenic peptides by class I Major Histocompatibility Complexes (MHC) and subsequent recognition of the peptide-MHC (pMHC) complex by cognate CD8+ T cells ^1,2^. Therefore, immunogenic peptides encompass attributes associated to two sets of features known as peptide-presentation and TCR recognition features ^3^. Among these, features attributed to MHC presentation have been shown to be more prominent compared to those attributed to TCR recognition. Examples include heavily conserved anchor positions and enriched motifs associated with distinct HLA types^4,5^. Indeed, recent cutting-edge models in predicting MHC presentation have shown impressive performance, exemplified by the widely used netMHCpan^6^ and other recently published models^7^.

The recognition features of peptides on the other hand are highly degenerate due to promiscuity of TCRs imposed by positive and negative selection to avoid immune blind spots^8^. In addition to peptides’ sequence-based recognition features, numerous other factors such as co-stimulation, proliferation and cytotoxicity underpin immunogenicity^1^. These factors collectively define the magnitude and shape of a T cell response and determine whether the response is elicited ^9^. Furthermore, nuances in various experimental assays evaluating T cell responses can produce noisy data, while the lack of a ‘true negative’ pool of peptides augments this complexity. Indeed, a ‘negative’ T cell assay only means a peptide failed to elicit a T cell response in a given experiment, perhaps due to the absence of a cognate T cell. This does not necessarily mean that a peptide is objectively non-immunogenic. Taken together, the identification of ‘immunogenic’ peptides has proven to be more challenging than identification of peptides presented by MHC molecules. Further challenges - e.g., limited numbers of known TCR-pMHC pairs - in identifying T cell antigens have recently been reviewed by Joglekar et al ^10^.

Click or tap here to enter text.Click or tap here to enter text.Despite these challenges, over the past decade several models have been presented to predict immunogenic peptides, leveraging different correlates of immunogenicity with varying levels of success. As we recently detailed^8^, a number of these studies have utilised sequence-based characteristics including amino acid features^13–17^, similarity to viral peptides^18^, sequence dissimilarity to self ^3,19^, association between peptide immunogenicity and their biophysical properties such as their structural and energy features^20^, as well as TCR recognition features^13,21^. Recently, Wells et al.^3^ comprehensively investigated a collection of parameters associated with neoantigen immunogenicity, grouped into 1) presentation features e.g., binding affinity and stability, hydrophobicity and tumor abundance and 2) recognition features e.g., agretopicity (the ratio of binding affinity between a mutated peptide and its wild-type counterpart), and foreignness (similarity of peptide of interest to previously characterised viral epitopes).

Despite the strong HLA-restriction of peptide recognition by conventional T cells, with most immunogenicity models, presentation features have not been deconvoluted from more subtle T cell recognition features in this manner, which - due to the prominence of MHC features in sequence data - may lead to models primarily predicting HLA ligands rather than immunogenic peptides. More recently though, studies such as those by Wells et al.^3^ and Schmidt et al.^22^ have shed light on this issue by disentangling features associated with presentation vs. those associated with T cell recognition ^3,22^.

In addition to the emerging consensus that predicting peptide immunogenicity involves deconvoluting MHC presentation and T cell recognition features, evidence is suggesting that different features will be required to predict immunogenicity of pathogenic epitopes vs. neoantigens, given fundamental differences in the mechanisms underpinning the respective T cell responses. Compared with pathogenic peptides, which are substantially different from the human proteome, neoantigens often exhibit only a single point mutation from the corresponding wild-type self-peptide^11^. This high sequence similarity between cancer neoantigens and self-peptides is likely subject to immune tolerance, thus focus has been placed on identifying features which permit neoepitopes to escape, which may be less applicable for pathogenic epitopes^12^.

Indeed, Richman et al. ^23^ and Devlin et al.^24^ have shown that dissimilarity to the self-proteome permits neoepitopes to escape from immune tolerance. However, we have recently shown that dissimilarity to self is limited in distinguishing immunogenic peptides from pathogens^12^, which indicate differences in the features required to predict immunogenic peptides from pathogens vs. cancer. Complicating matters further, existing models are primarily trained on pathogenic epitopes, some of which are used then to predict immunogenic neoantigens. Indeed, these important differences between pathogenic peptides and cancer neoantigens and their recognition by T cells indicate that separate features and training datasets would be required to reliably predict immunogenicity in these distinct settings.

Given this tapestry of complexity, it is unclear to what extent existing immunogenicity models can discriminate immunogenic peptides in human disease; although in mice, there is evidence that HLA ligand prediction alone may suffice in identifying such peptides. A recent study by Croft et al^25^ in vaccinia-virus infected C57BL/6 mice found that the majority (> 80%) of presented peptides by H-2D^b^ or H-2D^k^ are immunogenic, implying that HLA ligand predictors could be accurately used to identify immunogenic vaccinia-virus epitopes. This was evaluated in a recent study by Paul et al^26^., who benchmarked 17 models that are used primarily to predict antigen presentation. They showed each of these models can be used to predict immunogenic vaccina-virus peptides in mice at a relatively high level of confidence, albeit with some variation. They observed that netMHCpan v4.0 was the best performing model. However, we have recently observed that the fraction of presented peptides in humans which are immunogenic may be more variable (2.4% to 69% across different studies) than observed by Croft et al. in mice ^12^. It is unclear how much of this variation is a species-specific effect, or perhaps due to differences between experimental assays. Taken together, the extent to which immunogenicity models as well as antigen presentation features, can predict peptide immunogenicity in the context of different human disease settings, are open questions.

Numerous models over the past decade have been offered to predict peptide immunogenicity. These immunogenicity models have certainly contributed to our understanding of mechanisms underpinning T cell recognition in human disease, however, a systematic and unbiased evaluation of the performance of these models is not yet available and would provide a framework to guide identification of T cell targets in different immunological settings ^3,11,20^.

In this study, we evaluated and compared the performance of several publicly available models in predicting immunogenicity in human pathology settings. Given the crucial differences in- and features of-underlying mechanisms that invoke a T cell response, we illustrated the performance of these models in predicting immunogenic pathogenic epitopes and cancer neoantigens separately. In both settings, we found suboptimal model performance and considerable room for improvement. Furthermore, we explored several potential problems underpinning the suboptimal performance of some of these models and more generally the complexity of predicting peptide immunogenicity. We thus present a framework by which investigators can decide which immunogenicity classifier is more applicable to their data and research questions in the context of human disease.

## Results

### Evaluating model performance in predicting peptide immunogenicity

To perform an unbiased evaluation of existing models in predicting immunogenic peptides, we identified 7 publicly available models (see Methods for criteria details), each of which aim to predict whether an MHC-presented peptide may invoke a T cell response (i.e., whether a peptide is *immunogenic*). These models are named as the *IEDB model*^16^, *NetTepi*^15^, *iPred*^14^, *Repitope*^27^, *PRIME*^22^, *DeepImmuno*^17^ and *Gao*^28^ (Table 1). Additionally, given the observation by Paul et al.^29^ that netMHCpan 4.0 most accurately identified immunogenic vaccinia-virus T cell epitopes in mice, we included both eluted ligand (labelled *netMHCpan_EL*) and binding affinity (labelled *netMHCpan_BA*) outputs from netMHCpan 4.0 to assess their accuracy in a human setting. We therefore evaluated the performance of 9 models. We undertook the performance evaluation separately for pathogenic and cancer epitopes, due to inherent differences in these immunological settings.

**Table 1:** An overview of Immunogenicity Predictors which are evaluated in this study.

| <u>Model</u> | <u>Features and Calculations</u> | <u>Training Data</u> | <u>Language</u> | <u>Input Data</u> | <u>HLA Restriction</u> | <u>Reported Performance</u> |
| --- | --- | --- | --- | --- | --- | --- |
| IEDB Model<br>Calis, J. J. A. *et al.*<br>Properties of MHC Class I Presented Peptides That Enhance Immunogenicity. *PLoS Comput. Biol.* **9**, (2013). | The IEDB model captures sequence-based frequencies of specific amino acids to describe those which are more prevalent in immunogenic peptides compared with non-immunogenic peptides. Thus, ‘enrichment scores’ are combined with weights on amino acid positional importance for immunogenicity. The user is provided with a score between -1 and 1, to indicate likelihood of immunogenicity. | Data were compiled from the IEDB and three murine studies.<br>Strict data curation process excluded humans as a host for non-immunogenic peptides. Nine-mers were then selected and a redundancy filtering process was performed to avoid oversampling.<br>Dataset after redundancy filtering is not publicly available. | Python | Peptide(s) and HLA Allele | Yes. | ROC-AUC after 3-fold cross-validation: 0.65 |
| NetTepi<br>Trolle, T. & Nielsen, M. NetTepi: An integrated method for the prediction of T cell epitopes. *Immunogenetics* **66**, 449–456 (2014) | NetTepi was defined as the linear combination of binding affinity scores which were obtained by netMHCpan algorithm, binding stability scores which are obtained by netMHCstab (v1) and T cell propensity scores which are obtained similarly to the IEDB model:<br>$M = tTCellPropensity + sNetMHCstab + (1 - t - s)NetMHCcons$<br>Developers optimized weights $t$ and $s$, and defaults were given. *netMHCstab* and *netMHCcons* are called directly and canonically from NetTepi and individual scores from these algorithms are produced. These are integrated as above with the T cell propensity predictions. To our knowledge, the only difference between the T cell propensity component of NetTepi and the IEDB model, is that only default masking (for HLA-A0201) to encode for positional importance is employed by NetTepi, whereas for the IEDB model there are different settings based on chosen HLA. | T cell propensity component employed training data as above for the IEDB model. | Python | Peptides, HLA Allele, peptide length (8-14) | Yes, 13 HLA-A and HLA-B alleles | Average AUC0.1 values range between 0.9305 and 0.9652 |
| iPred<br>Pogorelyy, M. V. *et al.* Exploring the pre-immune landscape of antigen-specific T cells. *Genome Med.* **10**, 1–14 (2018) | iPred is a Multinomial Gaussian Process classifier, utilising expectation-maximisation and as input *iPred* takes peptide sequence(s) and computes 10 Kidera Features<sup>17</sup> averaged over all residues. The output of the model is a probability score reflecting the likelihood of T cell immunogenicity. | Chowell dataset was used to compute kidera factor vector sums for model training. | R | Peptide sequence | No | ROC-AUC: ~0.80 |
| Repitope<br>Ogishi, M. & Yotsuyanagi, H. Quantitative prediction of the landscape of T cell epitope immunogenicity in sequence space. *Front. Immunol.* **10**, (2019) | Repitope is a framework which probes public TCRs to discriminate immunogenicity. Repitope computes physiochemical properties based on mimicking the thermodynamics between pMHC and public TCR interactions. This framework termed ‘TCR-peptide Contact Potential Profiling’ (CPP) focuses on the TCRβ CDR3 sequences, which are primarily involved in the interactions with peptides presented onto MHC. After a highly dimensional feature (~6000) computation occurs, feature selection can be performed and then a probabilistic estimate of immunogenicity is computed for each epitope. Models can be extrapolated to sequences for which it was not trained to make predictions.<br>Repitope can compute probabilistic estimates of the immunogenicity for an original epitope and for all single amino acid variants, as well as the largest difference of immunogenicity between any variants to explore escape potential. Repitope also provides utility to predict CD4+ T cell immunogenicity for class II MHC. | Calis 2013 dataset, Chowell 2015 dataset, EPIMHC dataset, LANL HCV/HIV dataset, POPISK dataset, MHCBN dataset, TANTIGEN dataset. Please see following link for more details:<br>https://github.com/masato-ogishi/Repitope | R or Python. | Peptide sequences | No | ROC-AUC 0.76 |
| PRIME | PRIME is a predictor of immunogenic epitopes. The model captures and deconvolutes molecular properties of antigen-presentation and TCR recognition propensity. The authors were able to achieve this deconvolution by a ‘careful annotation and analysis of epitope residues with minimal impact on binding to HLA-I molecules’. PRIME is efficient and runs faster than many HLA-I ligand predictors. For the HLA binding component of the prediction, PRIME uses mixMHCpred. | Compiled from multiple studies. See table S1:<br>https://pubmed.ncbi.nlm.nih.gov/33665637/ | Python | Peptide sequences and a corresponding HLA allele for each peptide. | Yes | ROC-AUC >0.7, PR-AUC >0.10 < 0.16. |
| DeepImmuno | DeepImmuno-CNN is a convolution-neural network approach which predicts immunogenicity of pMHC complexes. A beta-binomial probabilistic model is fit to the training dataset in order to generate an immunogenicity score. Their immunogenicity score weights each pMHC complex based on the strength of available experimental evidence in the training dataset. A principal component analysis of 566 amino acid physicochemical properties encodes each amino acid sequence. | Initial training and validation occurred using data from the IEDB. Please see:<br>https://github.com/frankligy/DeepImmuno | Web portal or Python | Peptide sequence and corresponding HLA. Lengths 9/10 only | Yes | ROC-AUC 0.85 from cross-validation.<br>PR-AUC 0.81 from cross-validation. |
| Gao | A 'physics-based' learning model, aimed at predicting CTL epitopes. The model is trained and validated on peptides from HIV. | Peptide data from HIV patients. | R | Peptide and HLA allele. | Yes | ROC-AUC 0.66-0.71 |

**Table 2:**
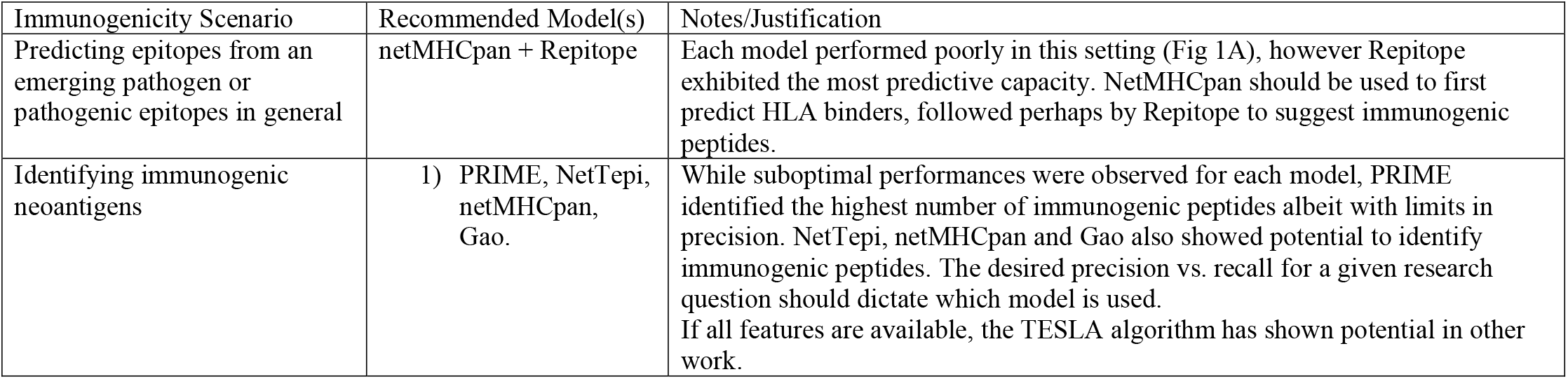
An overview of suggested applications for select immunogenicity predictors.

| Immunogenicity Scenario | Recommended Model(s) | Notes/Justification |
| --- | --- | --- |
| Predicting epitopes from an emerging pathogen or pathogenic epitopes in general | netMHCpan + Repitope | Each model performed poorly in this setting (Fig 1A), however Repitope exhibited the most predictive capacity. NetMHCpan should be used to first predict HLA binders, followed perhaps by Repitope to suggest immunogenic peptides. |
| Identifying immunogenic neoantigens | 1) PRIME, NetTepi, netMHCpan, Gao. | While suboptimal performances were observed for each model, PRIME identified the highest number of immunogenic peptides albeit with limits in precision. NetTepi, netMHCpan and Gao also showed potential to identify immunogenic peptides. The desired precision vs. recall for a given research question should dictate which model is used.<br>If all features are available, the TESLA algorithm has shown potential in other work. |

#### Evaluating model performance in predicting immunogenic pathogenic peptides

The ongoing efforts to understand T cell responses against SARS-CoV-2 have provided the community with an unprecedented number of functionally evaluated SARS-CoV-2 peptides. A unique and valuable characteristic of this data megapool is that – with the exception of several peptides from other coronaviruses (e.g., MERS, SARS-CoV) which share homology with SARS-CoV-2 ^30,31^ – many of these peptides have not been used in the training data of the models under investigation (Fig S1A) due to their recent identification. These peptides are therefore an ideal ‘test’ dataset for evaluating the performance of models in predicting immunogenic peptides from an emerging pathogen, enabling a benchmark of model performance in a realistic setting. Thus, we leveraged these data to shed light on the extent these models can be used to accurately identify immunogenic peptides upon the emergence of a novel pathogen.

To achieve this, we gathered all MHC class I SARS-CoV-2 peptides from the IEDB^32^ with T cell response information and supplemented the dataset with further peptides from VIPR^33^. Next, we applied two filters (see Methods for full details):

I. Length filter: We limited ourselves only on 9- and 10-mers because a) these lengths are the most common amongst CD8+ T cell targets and as a result the most prevalent lengths of class I peptides in our SARS-CoV-2 test data, b) these were the only lengths for which all 9 models are applicable.
II. HLA filter: We limited ourselves on the 13 HLA alleles for which all models were applicable (see Methods).

Identical homologs from e.g., MERS and SARS-CoV were retained (see Fig S1A), as upon the emergence of SARS-CoV-2, these conserved peptides may already have existed in model training data, therefore providing a more realistic testing scenario. Application of these filters left 878 SARS-CoV-2 peptides, of which ∼64% were immunogenic and ∼36% non-immunogenic. For added clarity, we assessed model performance using both receiver operating characteristic area-under-the-curve (ROC-AUC), which is commonly used in machine-learning contexts, and precision-recall area-under-the-curve (PR-AUC), which summarises model precision and recall and more accurately represents the balance of classes within the testing dataset.

The ROC-AUCs obtained from this task ranged from 0.504 for *DeepImmuno* to 0.574 for *Repitope* (Fig 1A), suggesting suboptimal performance for all models in identifying immunogenic epitopes from an emerging virus (Fig S1B, Supplementary Table 1). PR-AUCs ranged from 0.633 for *DeepImmuno* to 0.702 for *Repitope* (Fig 1B), which - given the proportion of immunogenic peptides in the dataset (∼64%) - suggests that select models may perform marginally better than random. Indeed, a follow-up bootstrap analysis (see Methods) revealed that the majority of models did not perform substantially better than the baseline using these data; although, with varying levels of significance, *Repitope* (ROC-AUC z-score = 4, PR=AUC z-score = 3.885) and the *IEDB model* (ROC-AUC z-score = 2.5, PR-AUC z-score = 2.659) did perform better than random (Fig S1C-D, Supplementary Table 2-3).

**Figure 1:**
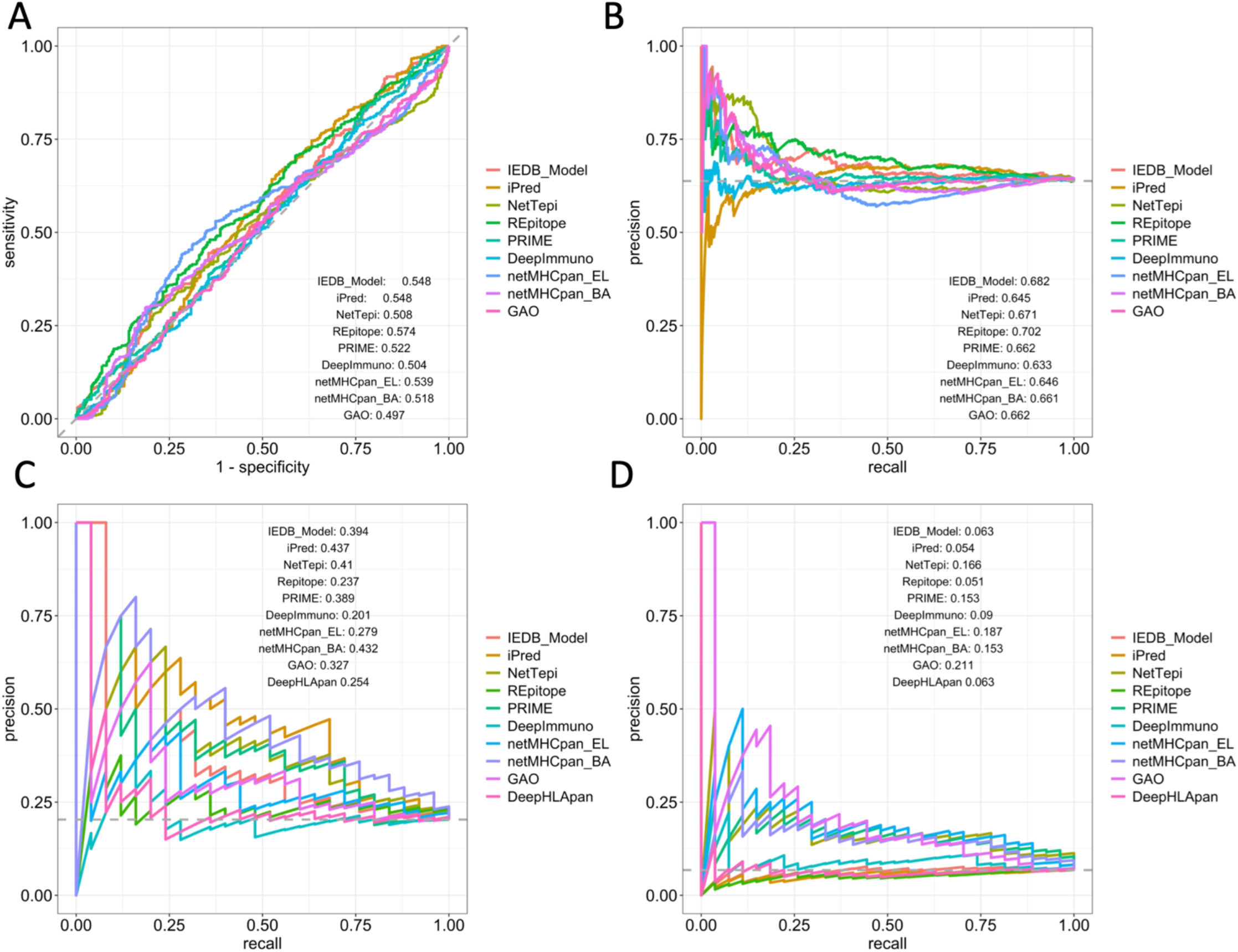
Models are not reliable in predicting epitopes for an emerging virus and exhibit room for improvement in predicting immunogenic cancer neoantigens. A) ROC-curves of models tested ‘as published’ against 681 SARS-CoV-2 peptides of lengths 9 and 10 (45% immunogenic). B) PR-curves of models tested ‘as published’ against the SARS-CoV-2 peptides. C) PR-curves showing model performance against the GBM neoantigen dataset. D) PR-curves showing model performance against the TESLA neoantigen dataset.

Taken together, neither the assessed HLA ligand predictors nor models specifically designed to predict immunogenicity, could reliably identify immunogenic peptides from an emerging pathogen. Thus, these data indicate that HLA ligand prediction is not sufficient to predict immunogenic epitopes from an emerging pathogen, suggesting that while presentation may be necessary it is not sufficient for T cell immunogenicity. Additionally, this analysis illustrates that current models which have been designed to incorporate further features which describe T cell recognition of pMHCI complexes, do not appear to outperform peptide presentation predictions in this setting. These insights therefore reveal a gap for an immunogenicity predictor to help extract immunogenic peptides from the pool of presented pathogenic peptides.

#### Evaluating model performance in predicting immunogenic cancer neoantigens

One key application for immunogenicity classifiers is to identify immunogenic cancer neoantigens that can activate CD8+ T cells for potential use as vaccine targets for personalised cancer immunotherapies^3,34^. Identifying immunogenic neoantigens is a ‘needle in a haystack’ problem, where one aims to find extremely small numbers of ‘positives’, from substantially imbalanced datasets. Indeed, multiple studies have observed that amongst predicted candidate cancer neoantigen datasets, validated *immunogenic* neoantigens comprise ∼6% ^3,35^, suggesting high false positive rates amongst current identification pipelines. Nevertheless, in this scenario, the ability of models to accurately identify small numbers of positives is paramount.

For highly imbalanced classification, ROC-AUC can be misleading as this metric can underrepresent the minority class^36^. Thus, we diagnosed model performance in predicting immunogenic cancer neoantigens using PR-AUC ^36^. We employed two independent cancer neoantigen datasets, both of which are intrinsically imbalanced: 1) our in-house glioblastoma (*GBM) dataset* and 2) a set of peptides gathered from the Tumor Neoantigen Selection Alliance (TESLA) consortium^3^ (see Methods for details on both datasets). For this cancer neoantigen setting, we have additionally evaluated the model ‘*DeepHLApan*,^37^, which is an immunogenicity predictor designed to identify cancer neoantigens.

Our *GBM* dataset comprises peptides which bind HLA-A*02:01 from glioblastoma cancer patients. We excluded any peptides observed in any model’s training data. The resulting dataset comprised peptides of lengths 9 and 10 (n=123), containing 25 (20%) confirmed immunogenic neoantigens. After testing these models against the *GBM* dataset, we observed suboptimal PR-AUCs, ranging from 0.20 for *DeepImmuno* to 0.437 for *iPred* (Fig 1C, Supplementary Table 4). With the exception of *DeepHLApan* which is hampered by false negatives, we observed a considerable number of false positives for each model (Fig S2A). Interestingly, *netMHCpan_BA, netMHCpan_EL* and *PRIME* identified the highest number (19, 18, 18 respectively) of the total 25 confirmed neoantigens (Fig S2A).

A bootstrap analysis revealed that despite suboptimal overall performance, the majority of models perform better than random (Fig S2B, Supplementary Table 5). For example, Z-scores evaluating deviation of the true PR-AUCs from a distribution of those achieved by random predictions demonstrate that *netMHCpan_BA* (z=5.2) possessed the most predictive power against this GBM dataset, followed by *iPred* (z=5.03) and *NetTepi* (z=4.38).

Next, we tested models against the publicly available ‘TESLA’ dataset. This dataset originally comprises cancer peptides amongst 13 class I alleles, of which we retained peptides experimentally tested against alleles for which all models are applicable, leaving peptides from 7 HLAs. Additionally, we excluded any peptides observed in any model’s training data. These filters resulted in 27 (∼6.7%) immunogenic and 372 (∼93%) non-immunogenic peptides of lengths 9 and 10.

We again observed suboptimal PR-AUC scores for each model, ranging from 0.051 for *Repitope* to 0.211 for *Gao* (Fig 1D, Fig S3A, Supplementary Table 6). After assessing predictive power as described previously, we observed that *Gao* (z=6.65), *netMHCpan_EL* (z=6.2), *NetTepi* (z=4.88), *PRIME* (z=4.55) and *netMHCpan_BA* (z=4.53) each performed better than random (Fig S3B, Supplementary Table 7). *Gao* utilises dissimilarity to self and similarity to viral peptides to compute immunogenicity of a peptide, which are features identified by Wells et al. as important in discriminating immunogenic neoantigens in the context of the TESLA dataset, which may explain *Gao’s* higher performance here compared with the previous scenarios. *PRIME* identified the highest number of TESLA neoantigens (26/27) followed by *netMHCpan_EL* which identified (22/27). Here, we observed high numbers of false positives for all models, including *DeepHLApan* (Fig S3A).

Overall, these data highlight the complexity of predicting cancer neoantigens. Despite contribution of these models to fostering our understanding of T cell recognition of neoantigens, these analyses additionally illustrate there exists considerable room for improvement in accurately identifying immunogenic neoantigens. It is of note that for most models, their performance is hampered by a considerable false positive rate, that contributes to low precision. It is also important to note that although in some medical settings, high false positive and low false negative rate is preferred, the ultimate aim of an immunogenicity classifier model is to achieve an optimal performance by balancing between false positive and false negative rates. Otherwise, models such as netMHCpan capable of accurate prediction of peptide processing and presentation for CD8+ T cells, would be sufficient to identify a target pool for further functional validation, as each *immunogenic* class I peptide should be presented. In this regard, the only model that exhibits a notable overall performance improvement compared with *netMHCpan* is the *Gao* predictor, although this was only observed against the TESLA dataset. We also observed cross-data inconsistency in performance of these models in predicting cancer neoantigens. Taken together, these results suggest that an avenue for improving the performance of these models beyond what can be achieved by HLA ligand predictors, is through improving the precision of these models to consistently identify the proportion of HLA ligands capable of invoking a T cell response.

In summary, we have illustrated suboptimal performance of existing models in predicting targets for T cell responses for both emerging pathogens and cancers. In what follows we therefore explore a few potential features contributing to difficulties of predicting T cell target peptides.

### Exploring potential underlying issues with model performances

Here, we aim to shed light on issues and inconsistencies in model performances, which we hope can guide avenues of research for future immunogenicity predictors. First, given differences in model performances across the previously examined scenarios, we explore differences in underlying features associated with immunogenicity between pathogenic vs. cancer settings, as well as potential reasons for contrasting performances within cancer settings. Secondly, due to HLA imbalances in training datasets and the low precision of the models, combined with their limited capacity to extend the performance achieved by HLA ligand predictor *netMHCpan*, we explore the extent to which these models may primarily predict antigen presentation.

#### Differences in discriminative features associated with immunogenicity for pathogenic vs. cancer peptides

Despite broad application of these models in predicting peptide targets for CD8+ T cell responses in both cancer and infection settings, there is no systematic comparative analysis of features associated to cancer vs. pathogenic peptide immunogenicity^12^. As discussed above in identifying immunogenic peptides in these two settings, we noticed inconsistent performances between cancer vs. pathogenic scenarios as well as *within* cancer settings. For example, the *Gao* model was superior against one of two cancer datasets, but performed poorly against pathogenic epitopes. These inconsistencies suggest potential differences in parameters leading to pathogenic vs. cancer neoantigen immunogenicity, which may translate into differences in the features required to discriminate immunogenicity in these two settings. We reasoned that such differences may contribute to the observed inconsistencies in model performances.

As mentioned, Richman et al.^23^ and Devlin et al.^24^ demonstrated that dissimilarity of a peptide to the self-proteome is associated with *neoantigen* immunogenicity. However, our recent work^12^ has demonstrated that dissimilarity to self is limited in discriminating immunogenic pathogenic peptides, suggesting that different features may be required to identify immunogenic peptides in pathogen vs cancer settings. Furthermore, given the wider availability of pathogenic compared with neoepitope datasets, immunogenicity predictors are often trained primarily on the former, which may further convolute predicting neoantigen immunogenicity.

For these reasons, we sought to investigate whether the observed variability in performances between pathogenic vs. cancer settings, may stem from differences in features attributed to immunogenicity. As discussed previously, Wells et al.^3^ comprehensively interrogated features associated with neoantigen immunogenicity, defining those related to antigen-presentation, or those related to T cell recognition. We therefore compared the capacity of these features previously associated with neoantigen immunogenicity, to discriminate immunogenic and non-immunogenic pathogenic peptides.

We first gathered class I associated pathogenic peptides from the IEDB (‘pathogenic’ dataset, see Methods), as well as the neoepitopes from Wells et al.^3^ (TESLA ‘cancer’ dataset). For this analysis we employed the entire TESLA dataset, comprising 608 peptides, of which ∼6% were immunogenic and ∼94% were non-immunogenic. The curated ‘pathogenic’ dataset consisted of 22,717 peptides in total, of which ∼25% were immunogenic and ∼75% were non-immunogenic. We then compared differences in the following presentation features: binding affinity and stability and the fraction of hydrophobic amino acids; the remaining features – agretopicity, foreignness and tumor abundance – are less applicable for pathogenic peptides.

We observed that cancer peptides may exhibit stronger MHC binding affinities than their pathogenic counterparts (Fig 2A-i). Indeed, for both non-immunogenic and immunogenic groups, cancer peptides possessed significantly stronger binding affinities compared with pathogenic peptides (2A-ii). By comparing binding affinities between immunogenicity status, we observed that immunogenic pathogenic peptides had stronger binding affinities than non-immunogenic pathogen peptides, which was consistent with Wells’ observations for cancer peptides (2A-iii). Interestingly, the median binding affinity for *non-immunogenic* cancer peptides is similar to that of *immunogenic* pathogen peptides (2A-iii), in particular beyond ∼100nM (2A-IV), suggesting that thresholds learnt from training data to discriminate immunogenicity may need to be tailored to pathogen or cancer settings. It is also important to note that, with the exception of HLA-C*07:02 (possibly due to limited numbers of peptides for this HLA), this trend was observed across a set of common HLAs (Fig S4A).

**Figure 2:**
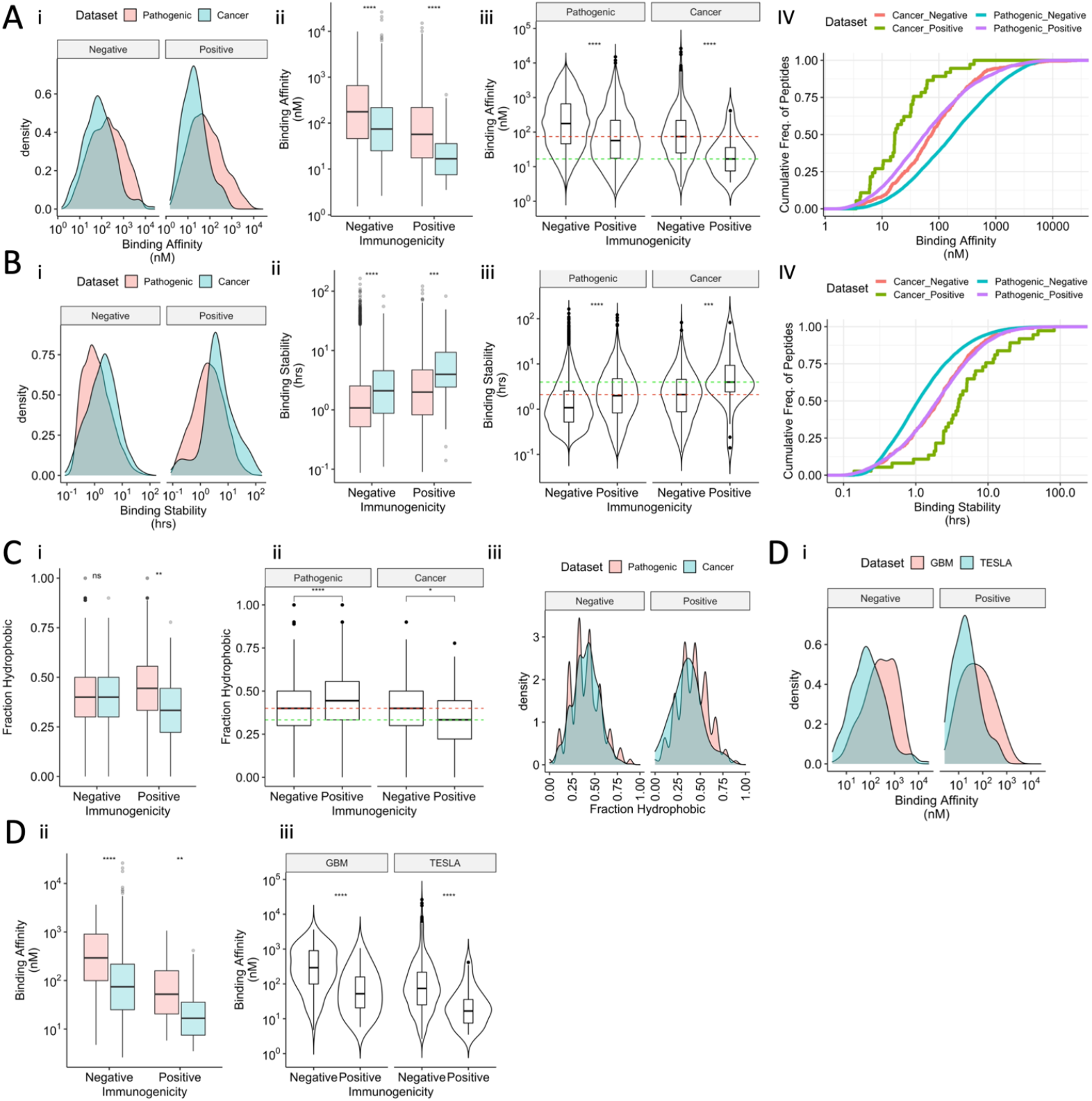
Differences in magnitude of discriminative antigen-presentation features associated with immunogenicity for pathogenic vs. cancer peptides: Analysis was performed examining A) binding affinities, B) binding stabilities, C) fraction of the peptide which is hydrophobic (fraction hydrophobicity) between pathogenic and cancer peptides. i) Density plots showing the distributions of binding affinities (Ai), binding stabilities (Bi) or fraction hydrophobicity (Ciii) for immunogenic and non-immunogenic peptides within pathogenic vs cancer datasets. ii) Boxplots comparing binding affinities (Aii), binding stabilities (Bii) or fraction hydrophobicity (Ci) of pathogen vs cancer peptides for both non-immunogenic and immunogenic peptides. iii) Violin plots comparing the binding affinities (A-iii), binding stabilities (B-iii), fraction hydrophobicity (C-ii) of immunogenic and non-immunogenic peptides, for both cancer and pathogenic datasets. Green and red dashed lines show the median of the binding affinities for the immunogenic and non-immunogenic cancer peptides respectively. IV) Line plots showing the empirical cumulative distributions of binding affinities (A-IV) and binding stabilities (B-IV), grouped by immunogenic or non-immunogenic within cancer or pathogenic peptide datasets. D-i) Density plots showing the distributions of binding affinities for immunogenic and non-immunogenic peptides between two independent cancer peptide datasets (GBM and TESLA). D-ii) Boxplots comparing binding affinities of GBM vs TESLA peptides for both non-immunogenic and immunogenic peptides. D-iii) Violin plots comparing the binding affinities of immunogenic and non-immunogenic peptides for GBM and TESLA cancer peptide datasets.

For MHC binding stability, the trends were consistent with those of binding affinity. First, cancer peptides exhibited greater stability compared to their pathogenic counterparts (Fig 2Bi – 2Bii). Similar to previous observations with binding affinity, we observed that while predicted binding stability can discriminate immunogenic and non-immunogenic pathogen peptides, the distributions of *non-immunogenic* cancer peptide stabilities are more comparable to those of *immunogenic* pathogenic peptide binding stabilities (Fig 2B-iii and Fig 2B-IV). These data provided further evidence that employment of the same thresholds or weights for MHC binding affinity or stability features, to discriminate immunogenic peptides in both pathogen and cancer settings, is likely to be ineffective. These observations of stronger binding affinities and more stable binding for cancer peptides may be due to facets of neoantigen identification pipelines, where one screens for high antigen presentation metrics to increase the likelihood of selecting better neoantigen candidates. This pre-selection for high presentation metrics may in turn contribute to skews in the datasets available for inference of MHC binding thresholds in discriminating immunogenic neoantigens.

As it comes to the ‘fraction of hydrophobicity’, we found it interesting that immunogenic pathogen peptides were more hydrophobic than their cancer counterparts (Fig 2C-i). Furthermore, this feature can discriminate immunogenic and non-immunogenic peptides in both pathogenic and cancer settings (Fig 2C-ii), although for pathogens there is a stronger trend, and in the reverse direction compared with cancer peptides. We additionally observed peaks around certain proportions for pathogenic peptides, and smoother distributions for their cancer counterparts (Fig 2C-iii). This is perhaps due to differences in the distributions of lengths between settings, where e.g., 65% of peptides are 9mers compared with 50% in cancer.

Next, as certain HLAs possess preferences for hydrophobic residues, we explored the capacity of this feature to discriminate immunogenic pathogen peptides across a set of common HLAs. Here, we observed variability in discriminating immunogenic peptides, suggesting this feature may need to be considered in an HLA-specific manner (Fig S4C). While these data suggest there is perhaps more weight for this feature in identifying immunogenic pathogen peptides vs. cancer – albeit in the reverse direction – further work is required to determine whether this is a biological characteristic or a technical artefact.

Inconsistencies in performances were also observed after testing the models against two independent cancer datasets. We therefore sought to examine whether immunogenicity parameters are likely to be linked to these inconsistencies. Consistent with the results comparing pathogens vs. cancer, we observed that binding affinity (Fig 2D-i, -ii, -iii), binding stability and fraction of hydrophobicity (Fig S4D) can discriminate immunogenicity with both cancer datasets (GBM and TESLA), while the thresholds required to discriminate immunogenicity are again likely to be different between the datasets. We were unable to explore the ‘recognition’ features associated with neoantigen immunogenicity as proposed by Wells, as our GBM dataset does not have tumor abundance information, which is a pre-requisite for the application of Wells’ recognition features. This task suggests that a fixed set of parameters may not be universal to all cancer datasets and that user input regarding the appropriate set of parameters, as well as interpretation of results, will be crucial in identifying immunogenic cancer neoantigens.

Overall, we have observed evidence for differences in the thresholds required for features to discriminate peptide immunogenicity in pathogens vs. cancer. Our observations indicate that a separate examination of presentation features and their weights in association with pathogen or cancer peptide immunogenicity is warranted. Furthermore, immunogenicity models are often trained primarily on pathogenic peptides. Therefore, as the distributions of binding affinities and/or stabilities for *immunogenic* pathogen peptides and *non-immunogenic* cancer peptides are comparable, this may in-part contribute to the substantial numbers of false positives generated by the models in identifying immunogenic cancer neoantigens, as well as inconsistencies in model performances between these immunological settings.

#### The effects of cross-HLA variation on predicting peptide immunogenicity

By exploring per HLA distributions of immunogenic and non-immunogenic peptides in the training data of these immunogenicity models, we observed substantial cross-HLA variation. Examples of these can be observed in *Repitope* (Fig S5A), the *IEDB model* (Fig S5B-C), and *iPred* (Fig S5D), where per HLA differences in numbers of positive (immunogenic) and negative (non-immunogenic) peptides are apparent. In fact, for *iPred* we observed that immunogenic peptides dominate the HLA-A supertype, whereas non-immunogenic peptides dominate HLA-B presented peptides in the training data (Fig S5D). As a result, HLA-A presenting peptides are more likely to receive a higher immunogenicity score from this model, regardless of their true immunogenicity status (Fig S5E).

We therefore sought to evaluate whether cross-HLA imbalances in immunogenic and non-immunogenic peptides may have contributed to suboptimal performances of some of the models. We hypothesised that without explicitly deconvoluting antigen-presentation features from TCR recognition propensity, training these models on HLA imbalanced datasets may lead to skewed predictions associated to HLA. This may in turn lead to predictions primarily of antigen presentation - due to more prominent MHC features ^5,7^ - rather than reflecting both peptide presentation and subsequent TCR recognition. We therefore aimed to explore whether the immunogenicity models (i.e., excluding HLA ligand predictors) exhibit skewed prediction scores toward dominant HLAs, and whether models thus predict prominent HLA types.

We first examined the distributions of the 10 (for consistency among models) most dominant HLAs in each model’s training data. We observed large skews toward and amongst HLA-A*02:01 for all models (Fig 3A). Indeed, by grouping peptides in each model’s training data by whether they bind HLA-A*02:01 (hereby A*02:01+) or not (A*02:01-), we observed considerable differences in the frequency of immunogenicity status per model training data (Fig 3B). We observed that with the exception of *IEDB* and *NetTepi*, non-immunogenic peptides dominate the A*02:01-group, meaning that non-immunogenic peptides are primarily presented by HLAs other than HLA-A*02:01 in model training data. It is plausible that this skewed distribution of prominent HLA features associated to immunogenicity status may lead to predictions skewed by HLA type.

**Figure 3:**
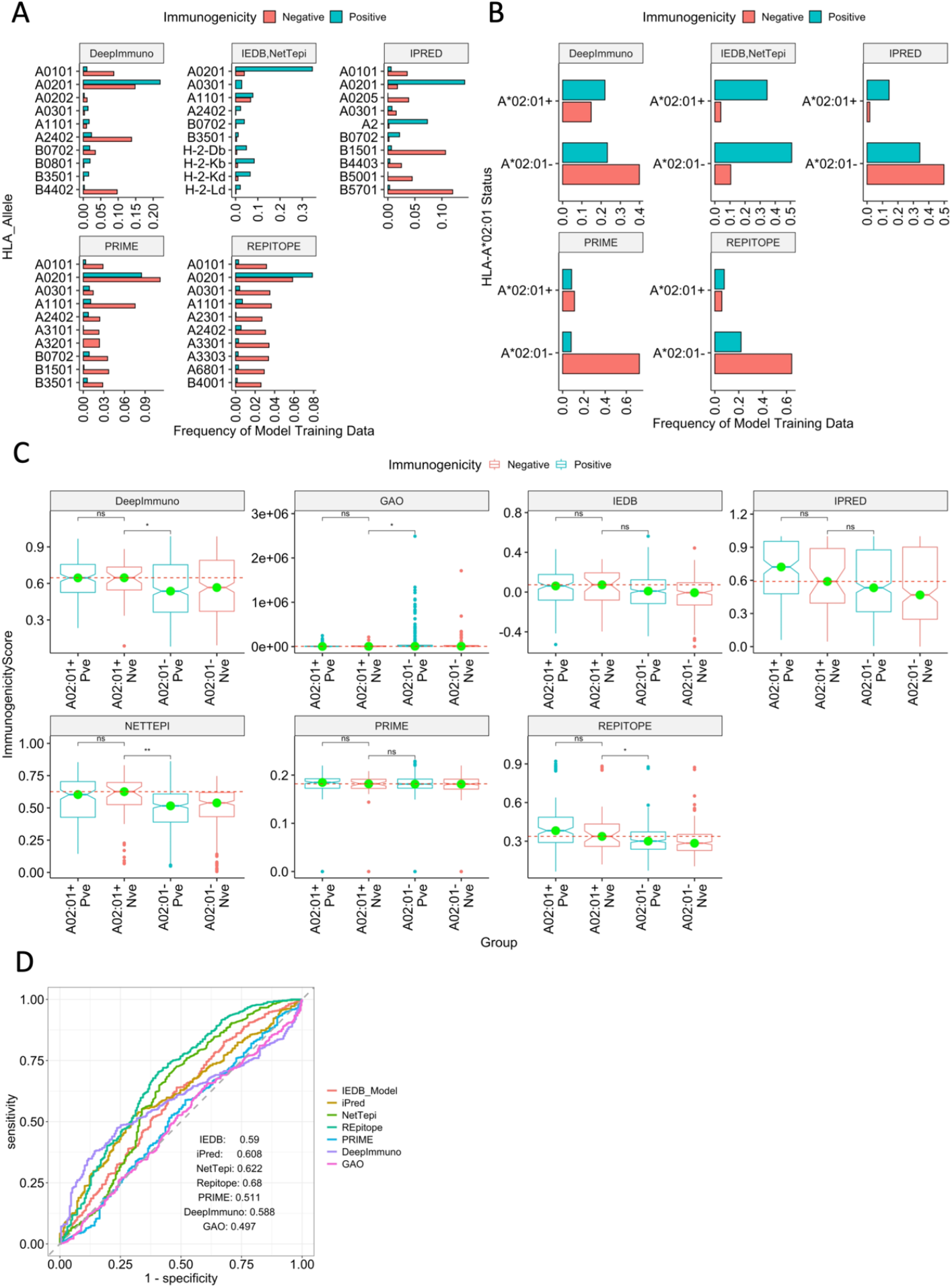
Effects of cross-HLA variation on predicting peptide immunogenicity. A) Barplots showing the distribution of immunogenic and non-immunogenic peptides for the top HLA alleles in each model’s training dataset. Gao’s training data is not included as the immunogenicity status are continuous. B) Barplots showing the distribution of immunogenic and non-immunogenic peptides for whether they bind HLA-A02:01 (labelled A*02:01+) or another allele (A*02:01-). C) Notched boxplots showing subtle differences between the immunogenicity scores created by each model, based on whether the peptide binds. HLA-A*02:01 (A02:01+) or another allele (A02:01-), and whether the peptide is immunogenic (Pve, labelled blue) or non-immunogenic (Nve, labelled red). D) ROC-AUCs of the models after taking scores produced to predict immunogenicity for SARS-CoV-2 peptides, but instead asked to predict HLA-A02:01 status.

We therefore hypothesised that these models may recognise prominent sequence features amongst HLA-A*02:01-binding peptides in test datasets, leading to inappropriately skewed model predictions. Therefore, we gathered the immunogenicity prediction scores generated by each model after evaluating their performance against our SARS-CoV-2 9- and 10-mer peptide dataset and grouped scores by whether the peptide is immunogenic or not, and whether it binds HLA-A*02:01 or not. First, for many examined models, we did not observe significant differences in immunogenicity scores between immunogenic and non-immunogenic peptides which bind HLA-A*02:01 (Fig 3C) (labelled A02:01+ Pve and A02:01+ Nve respectively). Furthermore, we found that *Repitope* (cohens-d=0.454), *DeepImmuno* (cohens-d=0.347), *NetTepi* (cohens-d=0.338), and to some extent *iPred* (cohens-d=0.206) predict *non-immunogenic* peptides that bind HLA-A*02:01 with higher scores than *immunogenic* peptides that bind remaining alleles; where on the contrary *immunogenic* peptides are expected to be predicted with higher immunogenicity scores. This analysis suggests that these models recognise dominant skewed HLA features associated with immunogenicity imbalances, which inappropriately skews model prediction scores, regardless of the true immunogenicity status of the peptide.

We next sought to analyse the extent to which these models detect peptide features associated with antigen presentation as opposed to T cell recognition. Here, we utilised prediction scores generated during the SARS-CoV-2 pathogenic epitopes analysis to instead predict whether a peptide engages HLA-A*02:01 or not (Fig 3D). Remarkably, we observed higher performance here for all models, compared with when tasked with predicting SARS-CoV-2 peptide *immunogenicity* (Fig 3D vs. Fig 1A), implying that for peptides from an emerging virus, all assessed immunogenicity predictors more capably predict dominant HLA features than T cell immunogenicity.

Overall, our data suggests that cross-HLA variation in the distribution of positive and negative peptides in model training data is highly likely to affect the peptide immunogenicity prediction, in a sense that some of these models in their existing settings predict HLA type more accurately than peptide immunogenicity. This insight suggests that suboptimal performance of these models is partly due to data limitations. Furthermore, our data suggest that future training datasets composed of more balanced immunogenic and non-immunogenic peptides for various HLA (or at least the common class I HLAs) are required for more accurate immunogenicity predictions. However, our work indicates that in the absence of such comprehensive datasets, the modelling strategies should consider how information extracted from HLA imbalances in training data affect model predictions and that future immunogenicity models should carefully model HLA restriction of T cell recognition.

## Discussion

In this study, we have demonstrated that despite great efforts, there is room for improvement to achieve accurate and reliable predictions of CD8+ T cells targets for an emerging virus such as SARS-CoV-2, or for a tumor of interest for the purpose of developing personalized (or stratified) treatments. We have additionally highlighted several issues underpinning suboptimal performances of these models with the hope of a) making potential users aware of these issues and guiding them towards strategies for use of an appropriate model for their tasks, and b) pointing future model developers to standing issues for further advancements and improvements.

For predicting CD8+ T cell targets for an emerging virus, these models do not seem to offer much improvement beyond MHC binding or presentation scores generated by models which predict HLA binding status such as *netMHCpan*, i.e., they may not offer much help extracting peptides that will trigger T cell response from the pool of presented peptides. Accurate and reliable prediction of CD8+ T cell targets would assist with several burning challenges, e.g., inference of escape mutations and/or the effects of mutations in CD8+ T cells targets on their immunogenicity.

For predicting immunogenic cancer neoantigens, we again illustrated that the assessed models only marginally outperform *netMHCpan. PRIME* and *NetTepi* were able to identify a high proportion of immunogenic neoantigens (high Recall), albeit at the expense of many false positives (low precision). Indeed, in most cases in this setting, the models produced many false positives. As immunogenic class I peptides must be presented by MHC, HLA ligand prediction with 100% accuracy could theoretically include 100% of immunogenic peptides (high recall), albeit with a high proportion of false positives (low precision) from those presented peptides which are not immunogenic. This concept is consistent with most observed model performances, and perhaps may in-part explain low precision observed with these models, as multiple predictors incorporate -and barely extend performance of-antigen presentation predictions.

Consensus is indeed emerging that presentation of a mutated peptide is insufficient for neoantigen immunogenicity ^3,11,35^. Our findings are consistent with this concept and indicate that additional features are required to consistently discriminate HLA ligands which can or cannot invoke T cell responses. Recently, the work of Wells et al. has provided a paradigm for such an approach. Consistent with Wells et al., our work suggests that parameters associated with neoantigen immunogenicity may require calibration for individual use cases. Further factors, e.g., differences between cancers, technical variations between experiments, and inherent human variation, are likely to compound this complexity.

Several recent studies have investigated some additional parameters associated with peptide immunogenicity that were not covered in the models that we have evaluated^3,11,20^. Riley et al.^20^ presented a structure-based approach, reporting increased performance against other models including the *IEDB model* and *NetTepi*. Despite this success, the authors suggest improvements to their model are necessary before wide adoption of their methodology. To our knowledge, their predictor is not yet publicly available, thus we were not able to evaluate its performance in the present study. Capietto et al. ^11^ recently supplied a framework for how mutation position contributes to neoantigen immunogenicity and proposed that the suboptimal landscape of neoantigen prediction, stems from a limited number of available tools which capture a variety of features associated with neoantigen immunogenicity. Indeed, future work should seek to examine the full spectrum of available parameters associated with neoantigen immunogenicity.

By investigating several potential factors contributing to suboptimal performance of these models in identification of immunogenic viral or cancer peptides, our work pointed towards both data- and model-associated issues. Data-associated issues include, i) small sample numbers especially for uncommon HLAs, an issue which is compounded for neoantigens, ii) cross-HLA imbalances of positive and negative peptides and iii) lack of true non-immunogenic peptides available in training datasets. Furthermore, the immunogenicity of a peptide is determined using different functional T cell response assays, which adds further noise to the data. With regard to cross-HLA imbalances, our findings suggest that skews in the distributions of immunogenic and non-immunogenic peptides per HLA in model training data may introduce bias into predictions. This observation is supported by Bassani-Sternberg et al^7^, where they showed that sequence similarity (i.e., HLA-I binding motifs) could effectively cluster peptides by respective HLA allele binding.

Model-associated issues suggested by our work include the use of universal parameter values and features for the identification of both viral and cancer antigens, training models on pathogenic peptides which are then used for prediction of cancer neoantigens and vice versa and limited consideration of HLA-restriction criteria of T cell recognition. Indeed, deconvoluting information extracted from anchor positions vs. TCR contact position, as opposed to modelling only full-length peptides appears to be an attractive approach for incorporating HLA-restriction, such as with work by Wells^3^ and Schmidt^22^. Furthermore, considering the lack of true negative training data, semi-supervised learning models e.g., generative variational autoencoder models seem appealing options.

Although in this study we sought to provide a general and an unbiased evaluation of the performances of existing immunogenicity models, there exist several limitations to our approach that are worth highlighting. First, we had to limit ourselves to specific HLAs and on peptide lengths (9 and 10) for which all models are applicable. More data from other HLAs and peptide lengths would be required for generalisation of our observations. Second, despite some potential benefits in re-training these models to further assess their reliability in different research settings, we chose to employ the models as trained by their authors. Key justifications for this are: a) the aim to provide a fair comparison across models, b) in most cases the end users would employ a model ‘as published’ and would not re-train them themselves. Additionally, retraining models may require recalibration of their parameters given new training datasets.

Third, for evaluating performance of models in identifying immunogenic peptides for emerging pathogens we performed this only - as a proof of concept - with a SARS-CoV-2 dataset that has not been used in training datasets of the models. Therefore, cross-pathogen variation in immunogenicity prediction is likely and highly important, but it is beyond the scope of this study. Finally, for our work regarding how HLA imbalances affect model predictions, we could only feasibly assess HLA-A*02:01 vs. remaining alleles, rather than HLA-A*02:01 vs. specific HLAs. More data from other HLAs will permit HLA-specific analyses.

The work of Croft et al.^25^ indicated that vaccinia-virus peptides presented by mice MHC molecules are highly translated into T cell recognition and response. It is therefore not surprising to see that in mice HLA ligand predictors e.g., netMHCpan 4.0 may accurately identify immunogenic epitopes ^26^. However, as we have observed in the present study and also previously reported^12^, in humans HLA presentation does not seem to be sufficient for T cell recognition. Therefore, additional features of peptide immunogenicity are required to assist extracting T cell targets from the pool of presented peptides in humans.

We envision that the insights provided in this study can assist end-users to make evidence-based decisions for which model and parameters to use with their data and research questions. Our work suggests that presentation features binding affinity, peptide stability and fraction of hydrophobicity are all associated with peptide immunogenicity for both cancer neoantigens and as well as viral peptides albeit with different parameter thresholds. While recognition features foreignness and agretopicity are associated with neoantigen immunogenicity, they are likely to be less applicable to viral peptides. Our work additionally suggests that for both cancer and pathogenic CD8+ T cell target identification, the first reliable filter would be their presentation status which is predicted by tools such as netMHCpan. For pathogenic peptides, this could then be followed by additional filters e.g., *Repitope*. For cancer neoepitopes, *PRIME* – which incorporates presentation predictions – could be employed although users should expect high levels of false positives.

Our work highlights an urgent need to improve immunogenicity predictions for emerging pathogens, as well as to improve the precision of models in identifying immunogenic neoantigens. We envision that addressing these concerns would create vast potential for highly accurate immunogenicity predictors, which could augment the efficiency of medical research and managing pandemic disease. We believe that our work will provide a useful and informative resource in the community, to help future users choose the right model and parameters for their tasks and also to help manage their expectations. We hope this study will assist future model developers in addressing issues highlighted here and also to guide design of future experiments to provide the required data.

## Methods

### Models selected for analysis

First, we gathered all publicly available models and excluded those which we were unable to perform comparative assessment. After exclusion (see Table S8), we were left with 7 models: the *IEDB model, NetTepi iPred, Repitope, PRIME, DeepImmuno* and *Gao*. netMHCpan 4.1 eluted ligand and binding affinity scores were additionally evaluated. The study reporting ‘PRIME’ focuses on predicting immunogenic neoantigens however the model is presented also as a predictor of pathogenic epitopes. All models with the exception of ‘*DeepImmuno’* were downloaded and ran locally. We were unable to run *DeerpImmuno* locally for unknown technical issues, therefore we instead used the webserver at https://deepimmuno.research.cchmc.org/

### Data Analysis

All analyses were performed in R 4.0.3. All visualisations (excluding ROC and PR curves) were produced using either the *ggpubr* or *ggplot2* packages. ROC curves were produced using the *pROC* package and PR curves were produced using the *yardstick* package. Confusion matrices and assessment metrics were computed using the *caret* package.

### Definition of immunogenic peptides

In the current study, we refer to immunogenic peptides as those which possess a ‘positive’ label in data repositories such as the IEDB. This label refers to peptides where a T cell response has been observed by e.g., an IFNγ ELISpot assay, although other techniques are commonly used.

### Model training data acquisition

We obtained the IEDB model training data from the supporting information of Calis et al 2013. This version of the data is prior to their redundancy filtering step and to our knowledge the final training data is not publicly available. The ‘Chowell’ dataset used to train iPred was obtained from the github repository hosting the classifier: https://github.com/antigenomics/ipred/tree/master/classifier

The *MHCI_Human* training data for the human class I Repitope model was obtained from the R package, hosted at https://github.com/masato-ogishi/Repitope

Training data for ‘PRIME’ was downloaded from the supplementary of the original publication^21^. Training data for ‘DeepImmuno’ was downloaded from https://github.com/frankligy/DeepImmuno#deepimmuno-cnn

### Performance Evaluation

All models were executed with default settings. For models where HLA restriction is considered, the corresponding HLA was provided to the model for each peptide of interest. For *NetTepi*, the length of the peptide is also provided at the command line.

For each model ROC curves were built using the *pROC* package and PR curves built using *yardstick*. To compute confusion matrices, binary classifications must be generated. Thus, from ROC curves, optimal threshold values for binary classification (Positive or Negative) were generated using the youden index. The youden index uses ROC curves to compute a threshold value which maximises the equation (1-sensitivity+specificity). For each model, the individual computed threshold value was used to classify prediction scores into ‘Positive’ or ‘Negative’ sequences and compiled in an additional ‘*ImmunogenicityPrediction’* column. For each model, confusion matrices were generated using the *confusionMatrix* function in the *caret* R package.

In each experiment – with the exception of SARS-CoV homologs in the SARS-CoV-2 experiment - peptides observed in model training data were excluded from performance evaluations. Model performance was evaluated through a combination of ROC-AUC, PR-AUC, precision and recall. Other metrics such as F1 score and Balanced Accuracy were also calculated. ROC-AUC curves show the performance of a model by perturbing thresholding and visualising the true positive rate (fraction of true positives / all true positives) against the false positive rate (fraction of false positives / all true negatives). Curve information is summarised using the area under the curve. Given a balanced dataset for binary classification (50% each classification), a random, unskilled model will have a ROC-AUC of 0.5, reflecting only the balance in the dataset. In contrast, a perfect model would have a ROC-AUC of 1.0. In a similar fashion, PR curves is a visualisation of model precision and recall (equations are described below) after perturbing thresholds. A perfect model would have a PR-AUC of 1.0, and a no skill classifier would reflect the balance in the data, i.e., using the above example, PR-AUC would be 0.5.

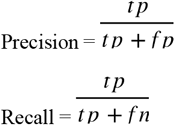

where ‘tp’ stands for ‘true positives’ and fp stands for ‘false negatives’.

### Evaluating model performance against SARS-CoV-2 peptides

The SARS-CoV-2 data were downloaded from the IEDB (accessed August 2021). Data were first filtered for class I binding peptides only (i.e., HLA allele information containing phrases *HLA-A, HLA-B*, or *HLA-C*). If contradicting pMHC observations (i.e., peptide *A*, HLA *y*, Immunogenicity both Positive and Negative in different experiments) were observed, the observation was assumed ‘positive’. NetTepi is only applicable for 13 HLA alleles (see below), therefore we only retained peptides assessed in the context of these alleles. We then filtered on 9 and 10mers, as *DeepImmuno* is only able to predict immunogenicity for these lengths. 49 SARS-CoV-2 peptides were observed in the collective model training data. After inspection, these peptides were from other coronaviruses. Given our aim to emulate a scenario with an emerging pathogen (where some peptides may have homology to other pathogens, but most are ‘unseen’), these were retained.

Each model was executed as published and prediction scores were generated for each peptide. ROC-AUC, PR-AUC, assessment metrics and confusion matrices were generated as described previously.

Alleles for which NetTepi is applicable: HLA-A*02:01, HLA-B*58:01, HLA-B*15:01, HLA-B*35:01, HLA-B*07:02, HLA-A*01:01, HLA-A*03:01, HLA-A*11:01, HLA-A*24:02, HLA-A*26:01, HLA-B*27:05, HLA-B*39:01, HLA-B*40:01.

### Bootstrap analysis to assess model predictive power

To assess predictive capacity of models against a test dataset, we first gathered the prediction scores generated by each model against the test dataset. These scores were not altered however, we randomly shuffled the immunogenicity label for each peptide. After each shuffling, we calculated PR-AUCs between the original model prediction scores and the newly shuffled immunogenicity labels. This process was repeated 1000 times, resulting in a distribution of ‘shuffled PR-AUCs’, reflecting performance of a distribution of random, unskilled models. We then compared the true ‘benchmarked PR-AUCs’ against this distribution of ‘shuffled PR-AUCs’, to provide a representation of how superior the model performance observed during benchmarking was to a distribution of unskilled models. Z-scores were calculated by subtracting the mean of the ‘shuffled PR-AUCs’ from the benchmarked PR-AUCs, and then dividing by the standard deviation of the ‘shuffled PR-AUCs’.

### Generating the ‘GBM’ dataset

These peptides were predicted using an in-house neoantigen identification pipeline and subsequently functionally validated. Prior to functional validation, these peptides were filtered to be of netMHCpan priority score > 50. In-vitro priming of healthy donors peripheral blood mononuclear cells (PBMC) and peptide stimulation of patient’s autologous PBMC was performed to find peptide specific T cells. Positive peptides were those where peptide specific T cells were identified by pMHC(HLA-A02:01)-tetramer staining. A peptide titration curve was then performed. To minimise variation in the current study, we additionally filtered for lengths 9- and 10-mer.

### Evaluating model performance against ‘GBM neoantigens’

Each model was executed as published and prediction scores were generated for each peptide in the ‘GBM’ dataset. ROC-AUC, PR-AUC, assessment metrics and confusion matrices were generated as described previously.

### Evaluating model performance against ‘TESLA consortium neoantigens’

The ‘TESLA’ dataset was downloaded from Wells et al^3^. In their study, 608 predicted neoantigens were derived from six patients. These predicted neoantigens were tested for immunogenicity, and 37 of them were found to be immunogenic. Here, test peptides were filtered for lengths (9 and 10) and HLA alleles (see above) for which all models are applicable

### Exploring differences in features associated with immunogenicity between pathogen and cancer datasets

The ‘TESLA’ dataset was acquired as above. All 608 peptides of lengths 8-14 were employed in this analysis. To curate the ‘pathogenic’ peptide dataset, we downloaded MHC class I peptides from the IEDB (accessed September 2021). This dataset was then supplemented with further peptides from the *Repitope* package’s data repository. We retained only peptides of lengths 8-14 and excluded any peptides from ‘antigen organisms’ containing the phrases “homo sapien” or “cancer”. Only peptides predicted to bind their corresponding HLA allele were retained for analysis. For analysis, netMHCpan4.0 was used to predict binding affinities of peptides to corresponding HLAs. netMHCstabpan was used to predict binding stabilities. The ‘fraction of hydrophobicity’ was calculated as the fraction of a peptide’s residues that were hydrophobic. Hydrophobic residues were considered as “V”, “I”, “L”, “F”, “M”, “W” and “C” ^3^.

### HLA imbalance in model predictions for pathogenic epitopes: Exploratory Analysis

We gathered the prediction scores generated by testing models against the SARS-CoV-2 dataset after being tasked with predicting immunogenicity. If a peptide was observed to bind HLA-A02:01 in this data, it was labelled as ‘HLA-A02:01+, while the remaining peptides were labelled ‘HLA-A02:01-.

### HLA imbalance in model predictions for pathogenic epitopes: Predicting HLA-A02:01+ peptides from SARS-CoV-2 peptide Immunogenicity Prediction Scores

Prediction scores generated by each model after the SARS-CoV-2 experiment to evaluate performance in predicting immunogenicity were gathered. Instead of producing ROC curves against the ‘Immunogenicity’ column however, ROC curves were produced against the binary classification of whether the peptide engages with HLA-A02:01 or not (i.e., HLA-A02:01+/-).

## Declarations

### Ethics Statement

N/A

### Consent for publication

N/A

### Availability of data and materials

Key datasets generated or analysed during this study are included in this published article [and its supplementary information files].

## Acknowledgements

N/A

## References

1. Zhang, N. & Bevan, M. J. CD8+ T Cells: Foot Soldiers of the Immune System. Immunity 35, 161–168 (2011).

2. Pennock, N. D. et al. T cell responses: naive to memory and everything in between. AJP: Advances in Physiology Education 37, 273–283 (2013).

3. Wells, D. K. et al. Key Parameters of Tumor Epitope Immunogenicity Revealed Through a Consortium Approach Improve Neoantigen Prediction. Cell 0, (2020).

4. Karnaukhov, V. et al. HLA binding of self-peptides is biased towards proteins with specific molecular functions. doi:10.1101/2021.02.16.431395.

5. Bassani-Sternberg, M. & Gfeller, D. Unsupervised HLA Peptidome Deconvolution Improves Ligand Prediction Accuracy and Predicts Cooperative Effects in Peptide– HLA Interactions. The Journal of Immunology 197, 2492–2499 (2016).

6. Reynisson, B., Alvarez, B., Paul, S., Peters, B. & Nielsen, M. NetMHCpan-4.1 and NetMHCIIpan-4.0: improved predictions of MHC antigen presentation by concurrent motif deconvolution and integration of MS MHC eluted ligand data. Nucleic acids research 48, W449–W454 (2020).

7. Bassani-Sternberg, M. et al. Deciphering HLA-I motifs across HLA peptidomes improves neo-antigen predictions and identifies allostery regulating HLA specificity. PLoS computational biology 13, e1005725 (2017).

8. Lee, C. H. et al. Predicting Cross-Reactivity and Antigen Specificity of T Cell Receptors. Frontiers in Immunology 11, 565096 (2020).

9. Paludan, S. R., Pradeu, T., Masters, S. L. & Mogensen, T. H. Constitutive immune mechanisms: mediators of host defence and immune regulation. Nature Reviews Immunology 1–14 (2020) doi:10.1038/s41577-020-0391-5.

10. Joglekar, A. V. & Li, G. T cell antigen discovery. Nature Methods 1–8 (2020) doi:10.1038/s41592-020-0867-z.

11. Capietto, A.-H. et al. Mutation position is an important determinant for predicting cancer neoantigens. (2020) doi:10.1084/jem.20190179.

12. H Lee, C., Antanaviciute, A., R Buckley, P., Simmons, A. & Koohy, H. To what extent does MHC binding translate to immunogenicity in humans? ImmunoInformatics 3–4, 100006 (2021).

13. Ogishi, M. & Yotsuyanagi, H. Quantitative prediction of the landscape of T cell epitope immunogenicity in sequence space. Frontiers in Immunology 10, (2019).

14. Pogorelyy, M. v. et al. Exploring the pre-immune landscape of antigen-specific T cells. Genome Medicine 10, 1–14 (2018).

15. Trolle, T. & Nielsen, M. NetTepi: An integrated method for the prediction of T cell epitopes. Immunogenetics 66, 449–456 (2014).

16. Calis, J. J. A. et al. Properties of MHC Class I Presented Peptides That Enhance Immunogenicity. PLoS Computational Biology 9, (2013).

17. Li, G., Iyer, B., Prasath, V. B. S., Ni, Y. & Salomonis, N. DeepImmuno: deep learning-empowered prediction and generation of immunogenic peptides for T-cell immunity. Briefings in Bioinformatics 00, 1–10 (2021).

18. Luksza, M. et al. A neoantigen fitness model predicts tumour response to checkpoint blockade immunotherapy. Nature 551, 517–520 (2017).

19. Bjerregaard, A. M. et al. An analysis of natural T cell responses to predicted tumor neoepitopes. Frontiers in Immunology 8, 1566 (2017).

20. Riley, T. P. et al. Structure based prediction of neoantigen immunogenicity. Frontiers in Immunology 10, (2019).

21. Schmidt, J. et al. Prediction of neo-epitope immunogenicity reveals TCR recognition determinants and provides insight into immunoediting. Cell Reports Medicine 2, 100194 (2021).

22. Schmidt, J. et al. Prediction of neo-epitope immunogenicity reveals TCR recognition determinants and provides insight into immunoediting. Cell Reports Medicine 2, 100194 (2021).

23. Richman, L. P., Vonderheide, R. H. & Rech, A. J. Neoantigen Dissimilarity to the Self-Proteome Predicts Immunogenicity and Response to Immune Checkpoint Blockade In Brief. Cell Systems 9, 375–382 (2019).

24. Devlin, J. R. et al. Structural dissimilarity from self drives neoepitope escape from immune tolerance. Nature Chemical Biology 16, 1269–1276 (2020).

25. Croft, N. P. et al. Most viral peptides displayed by class I MHC on infected cells are immunogenic. Proceedings of the National Academy of Sciences of the United States of America 116, 3112–3117 (2019).

26. Paul, S. et al. Benchmarking predictions of MHC class I restricted T cell epitopes in a comprehensively studied model system. PLoS Computational Biology 16, (2020).

27. Ogishi, M. & Yotsuyanagi, H. Quantitative prediction of the landscape of T cell epitope immunogenicity in sequence space. Frontiers in Immunology 10, 827 (2019).

28. Gao, A. et al. Predicting the Immunogenicity of T cell epitopes: From HIV to SARS-CoV-2. bioRxiv : the preprint server for biology (2020) doi:10.1101/2020.05.14.095885.

29. Paul, S. et al. Benchmarking predictions of MHC class I restricted T cell epitopes in a comprehensively studied model system. PLoS Computational Biology 16, e1007757 (2020).

30. Buckley, P. et al. HLA-dependent variation in SARS-CoV-2 CD8+ T cell cross-reactivity with human coronaviruses. bioRxiv 2021.07.17.452778 (2021).

31. Lee, C. H. et al. Potential CD8+ T Cell Cross-Reactivity Against SARS-CoV-2 Conferred by Other Coronavirus Strains. Frontiers in Immunology 11, 2878 (2020).

32. Dhanda, S. K. et al. IEDB-AR: immune epitope database - analysis resource in 2019. Nucleic Acids Research 47, W502–W506 (2019).

33. Pickett, B. E. et al. ViPR: an open bioinformatics database and analysis resource for virology research. doi:10.1093/nar/gkr859.

34. Roudko, V., Greenbaum, B. & Bhardwaj, N. Computational Prediction and Validation of Tumor-Associated Neoantigens. Frontiers in Immunology vol. 11 27 (2020).

35. Yadav, M. et al. Predicting immunogenic tumour mutations by combining mass spectrometry and exome sequencing. Nature 515, (2014).

36. Saito, T. & Rehmsmeier, M. The precision-recall plot is more informative than the ROC plot when evaluating binary classifiers on imbalanced datasets. PLoS ONE 10, (2015).

37. Wu, J. et al. DeepHLApan: A Deep Learning Approach for Neoantigen Prediction Considering Both HLA-Peptide Binding and Immunogenicity. Frontiers in Immunology 10, 2559 (2019).

