## Supplementary figures and images for "Evaluating performance of existing computational models in predicting CD8+ T cell pathogenic epitopes and cancer neoantigens"

### Supplemental Figure 1

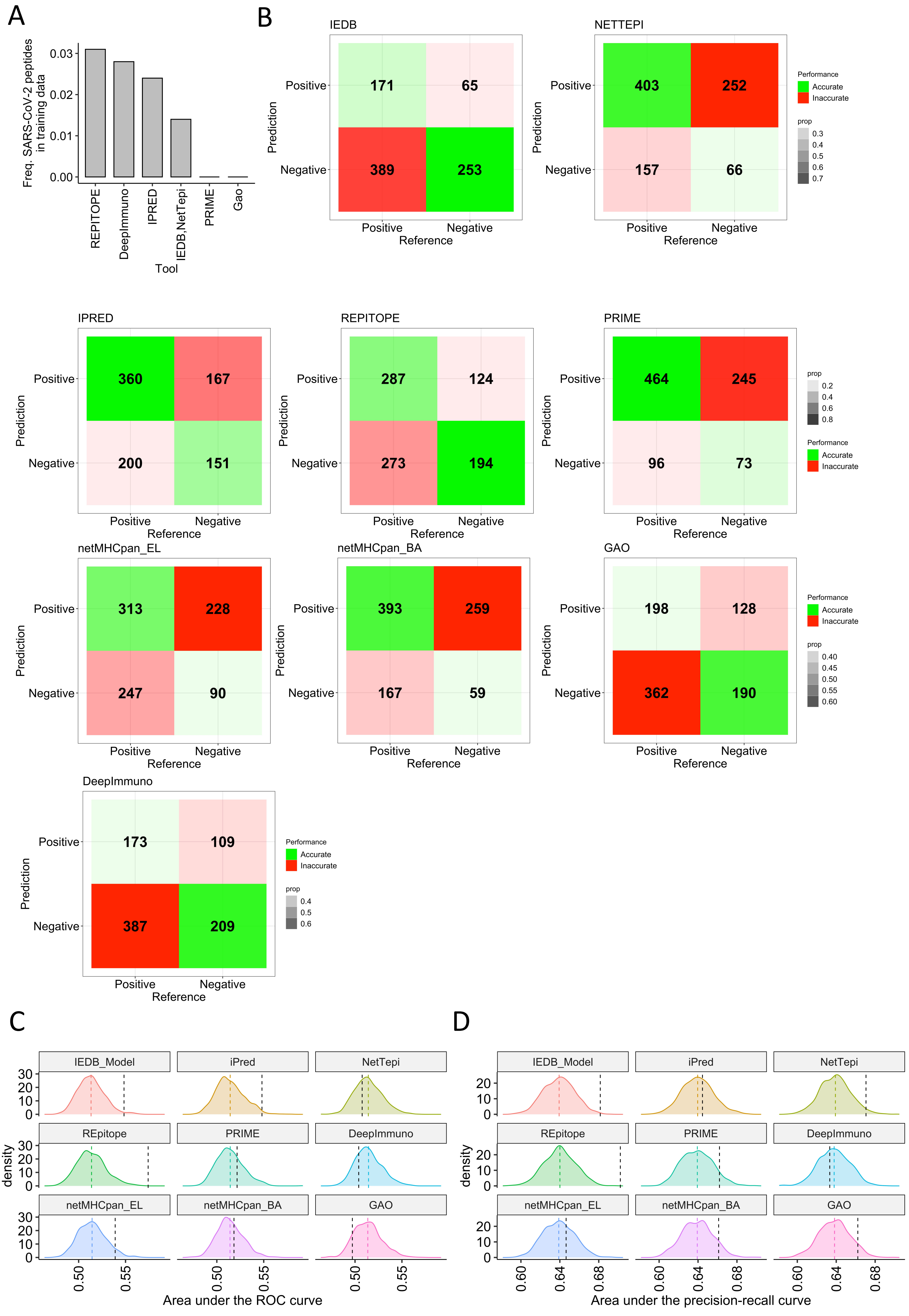

### Supplemental Figure 2

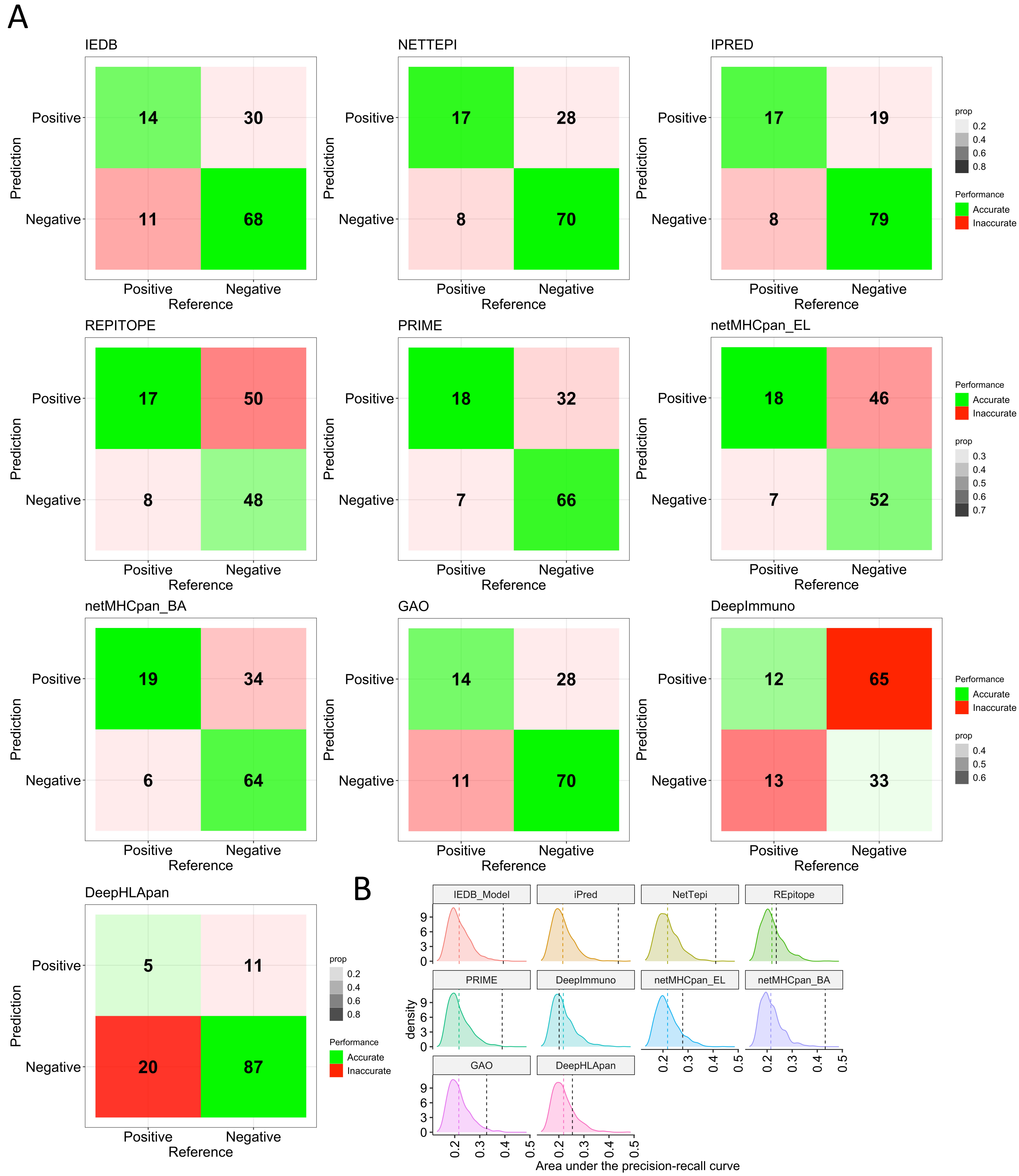

### Supplemental Figure 3

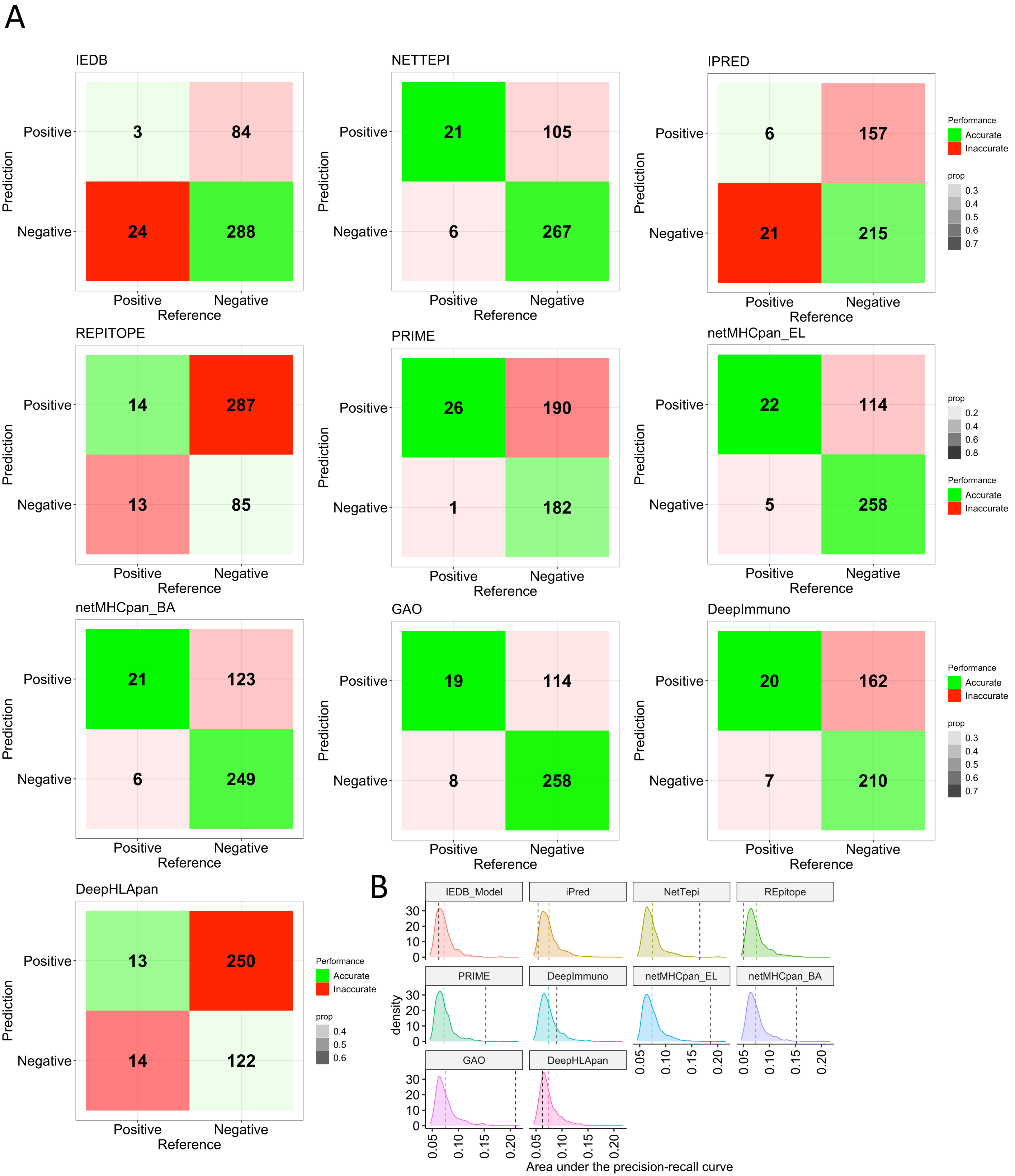

### Supplemental Figure 4

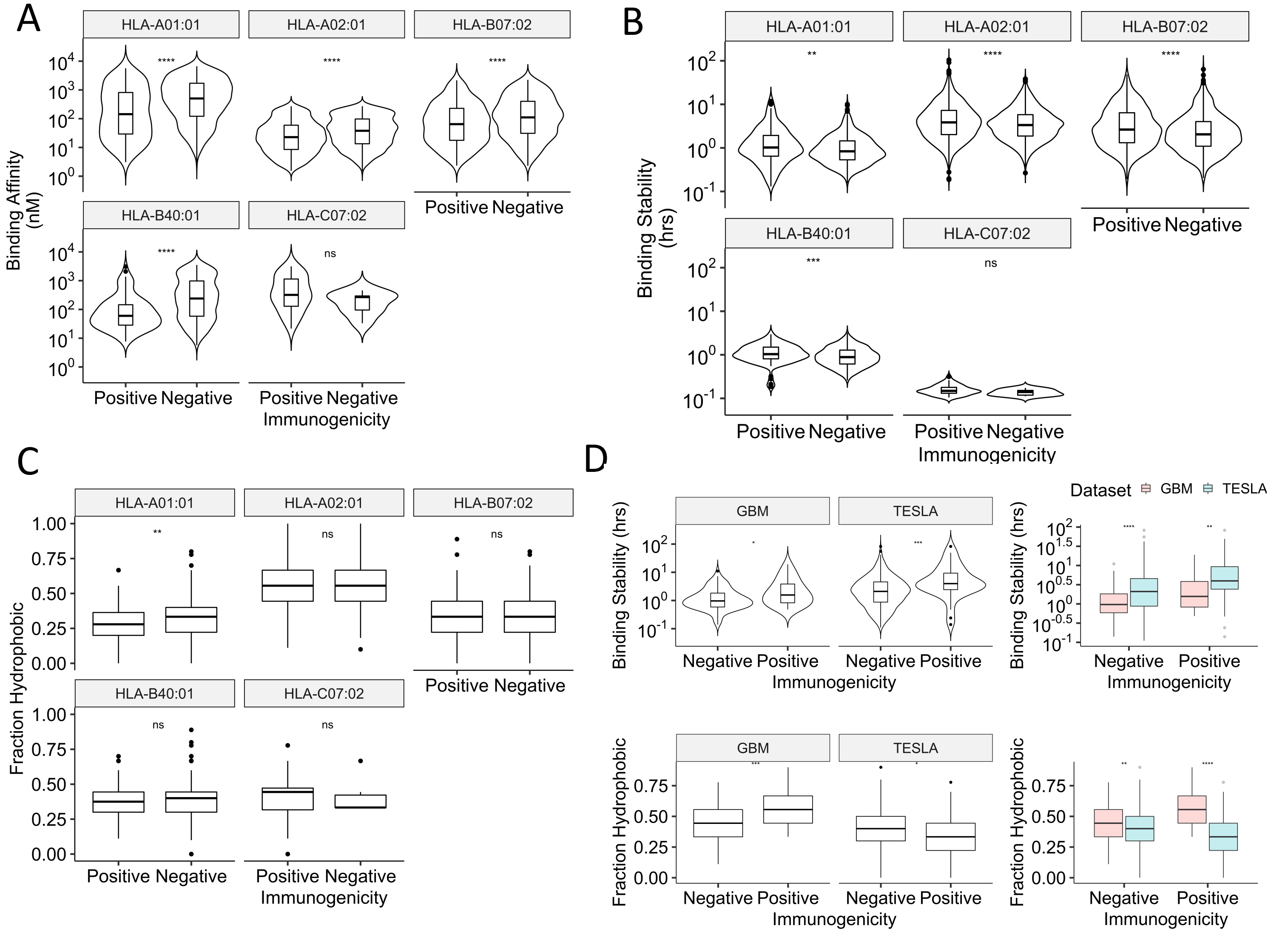

### Supplemental Figure 5

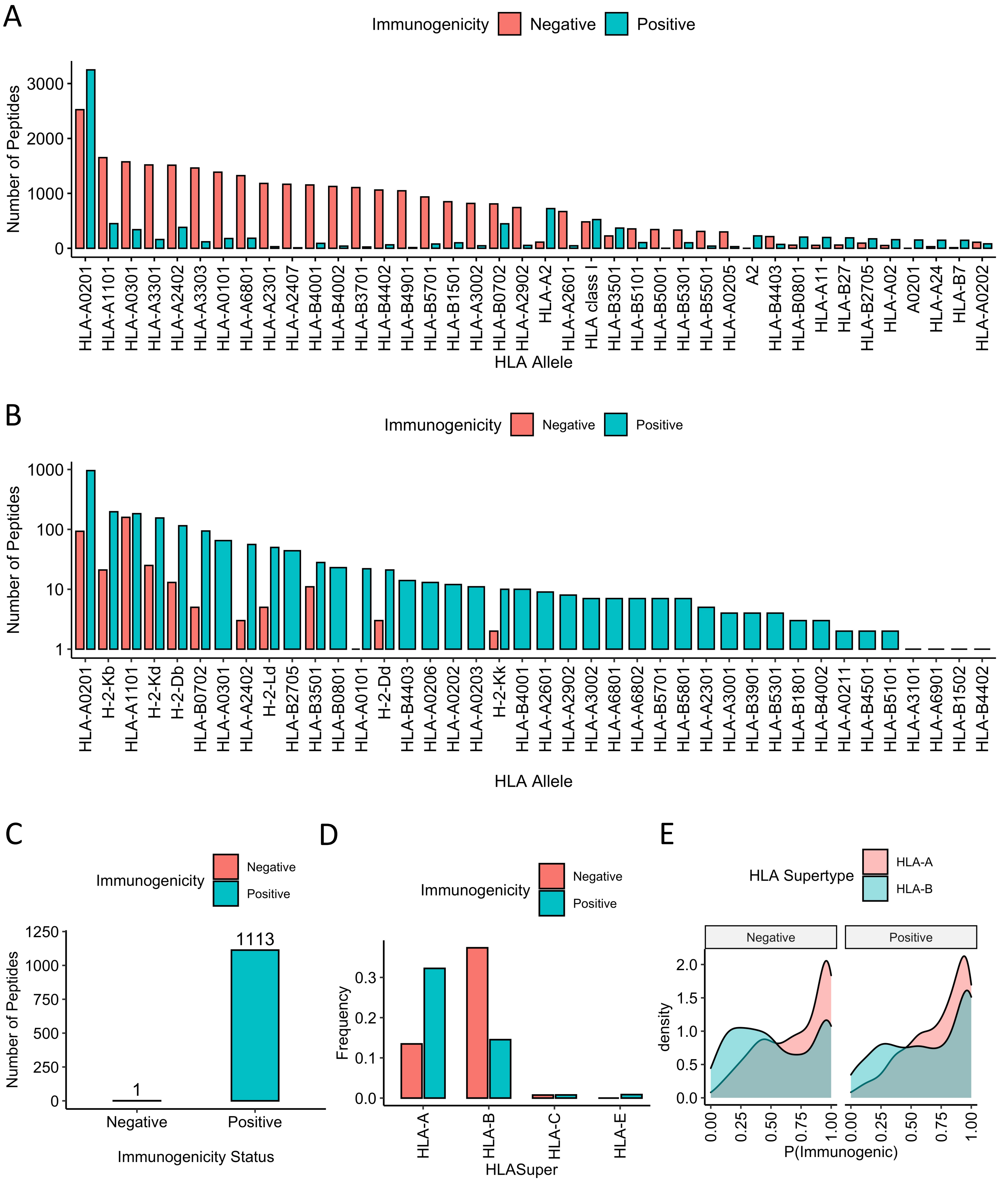
